# Multivalent Tau-Fyn interactions cooperatively arrest postsynaptic density condensate dynamics

**DOI:** 10.64898/2026.07.16.739052

**Authors:** Zheng Shen, Anne Bremer, Daxiao Sun, Maria-Sol Cima-Omori, Alf Honigmann, Stefan Becker, Markus Zweckstetter

## Abstract

Tau-Fyn signaling at postsynaptic sites is implicated in synaptic dysfunction and neuronal hyperexcitability in Alzheimer’s disease and related tauopathies, yet the molecular mechanisms by which Tau-Fyn interactions influence postsynaptic signaling assemblies remain poorly understood. Here, we reconstitute biomolecular condensates and membrane-associated assemblies that mimic the postsynaptic density (PSD) to investigate how Tau and Fyn regulate PSD organization and dynamics. We show that Tau and Fyn co-partition into PSD condensates and N-methyl D-aspartate receptor subtype 2B (NR2B)-associated PSD clusters, with both proteins preferentially enriching in the condensed phase relative to core PSD scaffold proteins. Enrichment of Fyn within PSD condensates is recapitulated by an SH3-SH2 fragment lacking the kinase domain, indicating that the kinase domain is dispensable for condensate partitioning. Fluorescence recovery after photobleaching reveals that Tau markedly reduces Fyn mobility and, together with Fyn, cooperatively drives a pronounced dynamic arrest of PSD condensate components, including the central scaffold protein PSD-95. Disruption of a major Tau-Fyn interaction hotspot by P216A/P219A mutations in Tau nearly completely restores the mobility of both Fyn and PSD-95, demonstrating that cooperative Tau-Fyn interactions are required for dynamic arrest. NMR spectroscopy identifies a multivalent interaction network between the Tau proline-rich region and the Fyn SH3 domain involving multiple binding motifs, with residues P216 and P219 acting as major interaction determinants. A 1.4 Å crystal structure of the Fyn SH3 domain bound to a Tau peptide reveals a ternary 2:1 Fyn:Tau assembly, providing structural support for multivalent Tau– Fyn engagement. Together, our findings establish a condensate-based mechanism by which multivalent Tau-Fyn interactions cooperatively regulate PSD condensate dynamics and provide a molecular framework linking Tau-Fyn signaling to synaptic dysfunction in tauopathies.

## Introduction

Synaptic dysfunction is an early and central feature of Alzheimer’s disease (AD) and related tauopathies, preceding overt neurodegeneration and strongly correlating with cognitive decline ^1, 2^. Increasing evidence indicates that Tau contributes directly to this process through functions that extend beyond its classical role as a microtubule-associated protein ^3, 4^. In addition to its axonal localization, Tau is present in dendritic compartments and postsynaptic structures, where it regulates neuronal excitability, glutamatergic signaling and synaptic plasticity ^5, 6^. Genetic and experimental studies have demonstrated that Tau is required for multiple pathological processes, including amyloid-β-induced synaptic deficits, network hyperexcitability and excitotoxic signaling ^7, 8, 9, 10^. Despite extensive evidence linking postsynaptic Tau to synaptic dysfunction, the molecular mechanisms through which Tau alters the organization and dynamics of postsynaptic signaling assemblies remain incompletely understood.

A central mediator of postsynaptic Tau function is the Src-family tyrosine kinase Fyn ^9, 11, 12^. Fyn acts as a key regulator of N-methyl-D-aspartate receptor (NMDAR) signaling by phosphorylating receptor subunits and associated signaling proteins and occupies a critical position within postsynaptic signaling networks ^13, 14, 15^. At excitatory synapses, Fyn interacts functionally with the scaffold protein PSD-95, which anchors NMDAR complexes within the postsynaptic density (PSD) and coordinates downstream signaling pathways ^16, 17, 18^. Multiple studies have demonstrated that Tau facilitates pathological Fyn-dependent signaling, and enhanced Tau-Fyn interactions have been implicated in Alzheimer’s disease, epilepsy and other neurological disorders ^19, 20, 21, 22^. Conversely, disruption of Tau-SH3 interactions ameliorates amyloid-β toxicity, synaptic dysfunction and neuronal hyperexcitability in several experimental systems ^9, 23^. Particularly notable is the observation that selective disruption of the sixth Tau PXXP motif, containing residues P216 and P219, is sufficient to rescue seizure susceptibility, network hyperexcitability and sleep abnormalities in mouse models, highlighting this interaction interface as a critical determinant of Tau-dependent pathology ^24^. Together, these findings identify Tau–Fyn interactions as a major pathological signaling node; however, the molecular mechanisms linking Tau–Fyn interactions to synaptic dysfunction remain poorly understood.

The PSD is a highly organized molecular assembly that concentrates neurotransmitter receptors, signaling enzymes and scaffold proteins beneath excitatory synapses ^25, 26^. Although traditionally viewed as a relatively static ultrastructural entity, recent work has revealed that the PSD is dynamically organized through biomolecular condensation ^27, 28, 29^. Core scaffold proteins, including PSD-95, GKAP, Shank and Homer, undergo liquid–liquid phase separation and assemble into condensates that recapitulate key structural and functional features of the PSD ^29^. These condensates provide a mechanistic framework for understanding how large signaling networks are assembled, maintained and remodeled within the confined environment of dendritic spines ^27, 28, 29^. Importantly, dynamic exchange of proteins within PSD condensates is thought to regulate receptor trafficking, signaling accessibility and activity-dependent remodeling of synaptic complexes ^30, 31, 32^. Changes in PSD dynamics can alter the mobility of scaffold and signaling proteins, thereby influencing the clustering, stability and activity of NMDAR signaling complexes and potentially affecting excitotoxic signaling pathways ^30, 31, 32^. Alterations in condensate dynamics and material properties are therefore emerging as potential mechanisms linking molecular organization of the PSD to both physiological and pathological regulation of synaptic function ^33, 34^.

Intrinsically disordered proteins are particularly well suited to regulate biomolecular condensates because they frequently engage in multivalent interactions capable of controlling condensate assembly, partitioning and dynamic behavior ^35, 36, 37^. Tau is a highly disordered protein containing multiple interaction motifs distributed throughout its proline-rich and repeat domains ^38, 39^. We recently demonstrated that Tau partitions into reconstituted PSD condensates through multivalent interactions with PSD-95 and markedly slows the dynamics of PSD proteins within these assemblies ^28^. These findings established Tau as an active regulator of PSD condensate dynamics rather than a passive client of the condensed phase and suggested a potential mechanism by which Tau could influence postsynaptic signaling ^28^. However, that work focused primarily on direct Tau-PSD-95 interactions and did not address how Tau cooperates with major postsynaptic signaling partners that are themselves strongly implicated in synaptic dysfunction. Given that PSD organization emerges from highly interconnected interaction networks rather than isolated binary interactions ^29, 34, 40^, additional Tau-binding partners may critically influence the dynamic behavior of PSD condensates.

Among these partners, Fyn represents a particularly compelling candidate. Tau contains multiple proline-rich PXXP motifs that interact with the Fyn SH3 domain, and residues P216 and P219 within the sixth PXXP motif have been identified as major determinants of Tau-Fyn binding ^41, 42, 43, 44^. More recently, structural and biophysical studies have revealed that Tau engages the Fyn SH3 domain through multiple alternative binding sites distributed across the proline-rich region, suggesting a dynamic and multivalent interaction network rather than a single canonical interaction ^41^. Such multivalent interactions are a defining feature of biomolecular condensates and frequently generate emergent collective behaviors that cannot be predicted from individual binary interactions alone. These observations raise the possibility that multivalent Tau-Fyn interactions may cooperatively regulate PSD condensate dynamics and thereby influence the organization of postsynaptic signaling assemblies. Whether such interactions contribute to the enrichment, organization and dynamic behavior of Fyn within PSD condensates, alter the mobility of PSD scaffold proteins, or provide a mechanistic basis for condensate remodeling has remained unknown.

Here, we sought to determine how Tau-Fyn interactions influence the organization and dynamics of PSD-like molecular assemblies. To address this question, we combined reconstituted PSD condensates and membrane-associated PSD clusters with quantitative partitioning analysis, fluorescence recovery after photobleaching, NMR spectroscopy and X-ray crystallography. Using complementary biophysical and structural approaches, we investigated whether multivalent Tau-Fyn interactions regulate the dynamic behavior of PSD condensates and defined the molecular basis of this regulation.

## Results

### Tau and Fyn partition together into PSD-like liquid droplets and membrane-associated clusters

We previously demonstrated that Tau partitions into reconstituted PSD condensates through multivalent interactions with PSD-95 and modulates condensate dynamics ^28^. However, PSD organization emerges from highly interconnected interaction networks rather than isolated protein interactions, raising the possibility that additional Tau-binding partners contribute to the regulation of PSD condensates. Because local enrichment of interacting proteins is a prerequisite for condensate-mediated regulation, we first asked whether Tau and the Src-family kinase Fyn partition into the same PSD-like assemblies (Fig. 1A).

**Fig. 1.**
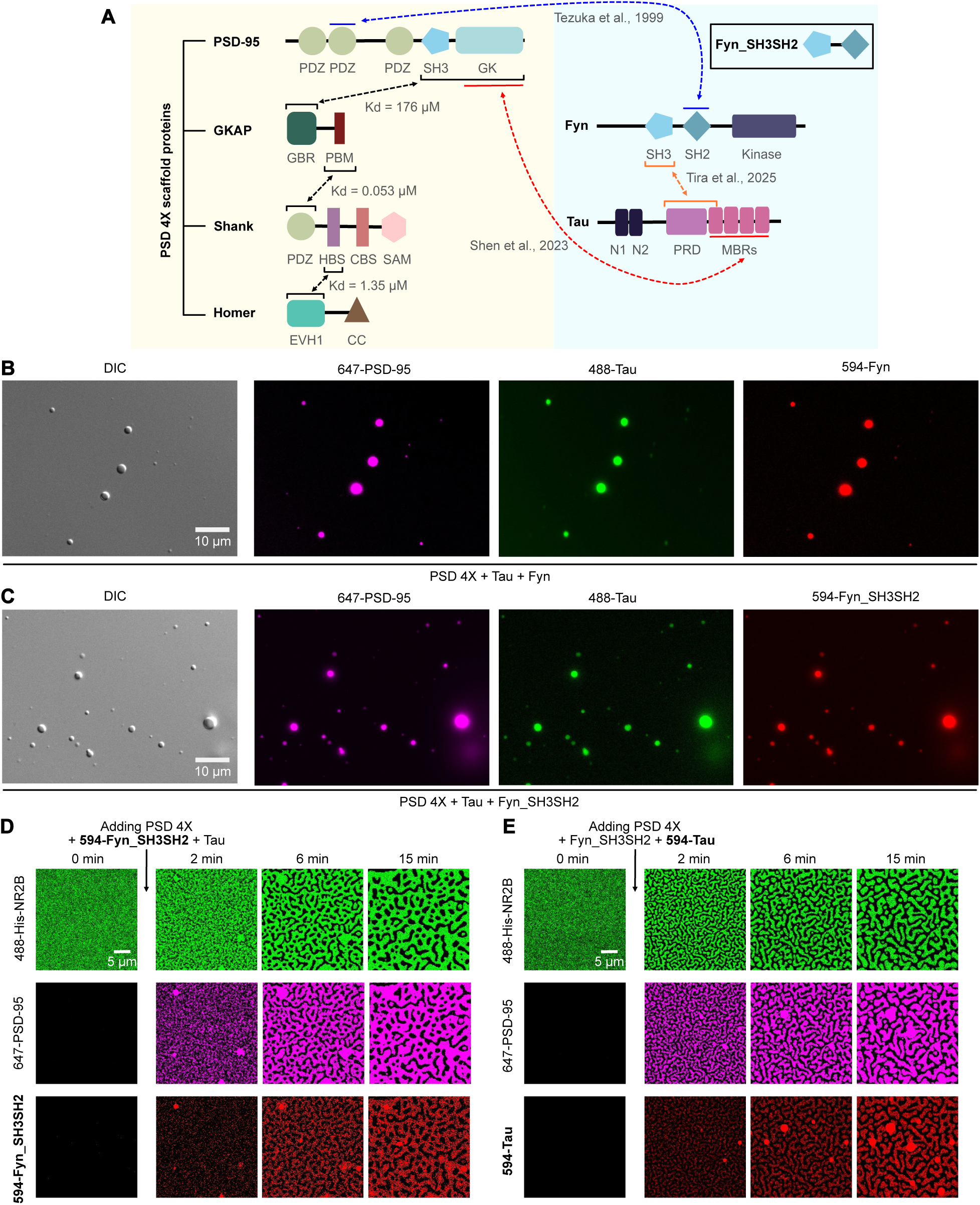
Tau and Fyn co-partition into PSD condensates and membrane-associated PSD clusters. **A.** Schematic diagram illustrating the major protein components used in this project: PSD-95, GKAP, Shank, Homer, full-length 2N4R Tau and Fyn. Black dotted arrows indicate the protein-protein interactions among the receptors and four PSD scaffold proteins, with binding affinities reported previously ^29^. The blue dotted arrow indicates the previously reported interaction between the PDZ2 domain of PSD-95 and the SH2 domain of Fyn ^16^. The red dotted arrow indicates the interaction between Tau and PSD-95 ^28^. The orange dotted arrow indicates the interaction of the Fyn SH3 domain with the proline-rich domain and R1 region of Tau ^41^. **B, C.** DIC and fluorescent images of PSD 4X condensates (containing PSD-95/GKAP/Shank/Homer) with Tau and Fyn proteins. Full-length Fyn was used in (B) and Fyn_SH3SH2 was used in (C). PSD-95, Tau and Fyn proteins were labeled with Alexa 647, Alexa 488 and Alexa 594, respectively. The concentration of each protein was 5 µM. **D, E.** Time-lapse confocal microscopy of Alexa 488-labeled His-NR2B clustering on supported membranes after adding four major PSD scaffold proteins (PSD 4X), Fyn_SH3SH2, and Tau. Alexa 647 labeled PSD-95 was used in both (D) and (E), Alexa 594-Fyn_SH3SH2 was used in (D) and Alexa 594-Tau was used in (E).

To investigate the partitioning of Tau and Fyn into PSD-like condensates in solution, we mixed the four PSD scaffold proteins PSD-95, GKAP, Shank and Homer (PSD 4X) with full-length human 2N4R Tau and full-length Fyn at an equimolar concentration (Fig. 1B). Phase-separated droplets formed immediately upon mixing (Fig. 1B). Fluorescent microscopy revealed strong colocalization of PSD-95, Tau and Fyn within the condensed phase, indicating that both Tau and Fyn partition into PSD condensates (Fig. 1B).

Fyn contains SH3, SH2 and kinase domains, and previous studies have implicated SH3-mediated interactions in Tau binding ^41, 42^. To determine whether the kinase domain contributes to condensate partitioning, we generated a truncated Fyn construct comprising only the SH3 and SH2 domains (Fyn_SH3SH2). When mixed with PSD 4X scaffold proteins and Tau, Fyn_SH3SH2 partitioned into PSD condensates together with PSD-95 and Tau, similarly to full-length Fyn (Fig. 1C). These findings indicate that the kinase domain is dispensable for partitioning into PSD condensates and suggest that the SH3–SH2 region is sufficient to support Tau/Fyn co-enrichment within the condensed phase.

We next investigated whether Tau and Fyn also partition into membrane-associated PSD assemblies. To reconstitute membrane-associated clusters, PSD components were assembled on supported lipid bilayers containing membrane-tethered His-NR2B, a construct designed to mimic the tetrameric membrane-associated organization of NMDA receptors by fusing the last five amino acids of NR2B to the C-terminus of the tetrameric GCN4 coiled-coil domain ^28, 29^. Because Tau and Fyn_SH3SH2 were labeled with the same fluorophore (Alexa Fluor 594), two parallel experiments were performed under identical conditions, one containing labeled Fyn_SH3SH2 (Fig. 1D) and the other containing labeled Tau (Fig. 1E). Following addition of PSD components together with Tau and Fyn_SH3SH2, His-NR2B rapidly formed membrane-associated clusters within 15 min. Fluorescence microscopy demonstrated extensive overlap between Fyn_SH3SH2, PSD-95 and His-NR2B signals (Fig. 1D), indicating incorporation of Fyn_SH3SH2 into membrane-associated PSD assemblies. Similarly, Tau fluorescence colocalized with PSD-95 and His-NR2B clusters (Fig. 1E), demonstrating incorporation of Tau into the same assemblies.

Together, these results show that both Tau and Fyn partition into PSD-like condensates in solution as well as membrane-associated PSD clusters. The co-enrichment of Tau and Fyn within the same assemblies establishes the physical basis for potential cooperative interactions and suggests that multivalent Tau-Fyn interactions could contribute to the regulation of PSD condensate organization and dynamics.

### Disease-relevant Tau-Fyn interactions promote dynamic arrest within PSD-like condensates

Because local enrichment of interacting proteins is a prerequisite for condensate-mediated regulation, we next investigated the partitioning behavior and dynamic properties of Tau and Fyn within PSD-like condensates. Previous studies have shown that disease-associated Tau reduces Fyn mobility within hippocampal dendritic spines, suggesting that Tau–Fyn interactions may influence the dynamic behavior of postsynaptic signaling complexes ^45^. We therefore asked whether Tau and Fyn exhibit altered mobility within reconstituted PSD condensates and whether disease-relevant Tau-Fyn interaction motifs contribute to this process.

To minimize indirect contributions from additional PSD scaffold proteins, we reconstituted condensates using only PSD-95 and GKAP (PSD 2X) in the presence of full-length human 2N4R Tau and Fyn_SH3SH2 at equimolar concentrations (Fig. 2A). Differential interference contrast (DIC) microscopy revealed rapid formation of phase-separated droplets upon mixing, and fluorescence microscopy confirmed partitioning of both Tau and Fyn_SH3SH2 into the condensed phase (Fig. 2A).

**Fig. 2.**
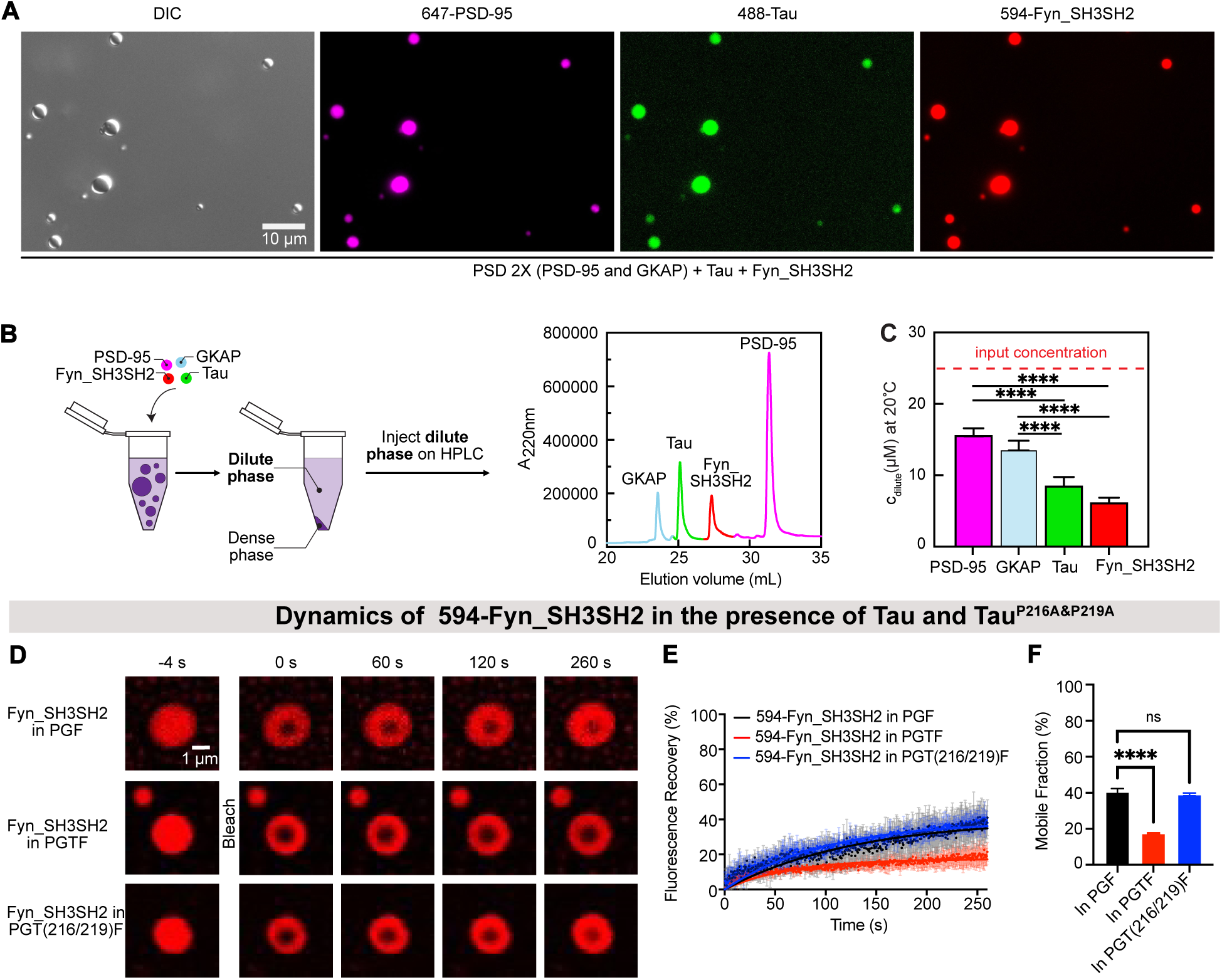
Disease-relevant Tau-Fyn interactions arrest Fyn dynamics in PSD condensates. **A.** DIC and fluorescent images of PSD 2X (PSD-95 and GKAP) condensates with Tau and Fyn_SH3SH2. PSD-95, Tau and Fyn_SH3SH2 were labeled with Alexa 647, Alexa 488 and Alexa 594, respectively. 25 µM of each protein was used. **B.** Schematic of the approach used to quantify the protein concentration of PSD-95, GKAP, Tau and Fyn_SH3SH2 in the dilute phase of PGTF condensates by analytical HPLC ^66^. Phase separation was induced by mixing the proteins at an equal molar ratio (25 μM each), followed by centrifugation to separate the dense and dilute phases. The protein concentrations in the dilute phase using 220 nm detection channel. **C.** Protein concentrations in the dilute phase (c_dilute_) of PSD-95, GKAP, Tau and Fyn_SH3SH2. Individual data points (n = 4) are shown, with error bars representing +/- SD; ordinary one-way ANOVA test: ****p < 0.0001. **D.** Representative FRAP images of Alexa 594-labeled Fyn_SH3SH2 in PSD-95/GKAP/Fyn_SH3SH2 condensates without (“PGF”; top), with Tau (“PGTF”; middle), and with Tau^P216A/P219A^ (“PGT(216/219)F”; bottom). **E.** FRAP-based quantification of 594-Fyn_SH3SH2 dynamics. Data are presented as mean values +/- SD from n independent experiments. The number of experiments (n) for Fyn_SH3SH2 in PGF, PGTF and PGT(216/219)F condensates are 5, 4 and 3, respectively. The FRAP curves (dots) were fitted with a mono-exponential function (solid-lines). **F.** Mobile fractions of Fyn_SH3SH2 were derived from the fitted averaged FRAP curves, which were averaged from n independent experiments. For each condition, n corresponds to the number of independent FRAP experiments indicated in (E). Error bars represent standard deviation of curve fits; unpaired and two-tailed t-test: ****p ≤ 0.0001.

To quantitatively assess partitioning, we separated the dilute and dense phases by centrifugation and measured protein concentrations in the dilute phase using analytical high-performance liquid chromatography (HPLC) (Fig. 2B). All components were initially present at equal concentrations (25 μM each). Quantitative analysis revealed that PSD-95 (15.7 μM) and GKAP (13.7 μM) remained substantially more abundant in the dilute phase than Tau (8.7 μM) and Fyn_SH3SH2 (6.3 μM) (Fig. 2C). These results demonstrate that Tau and Fyn_SH3SH2 preferentially partition into the condensed phase relative to the core PSD scaffold proteins, resulting in strong local enrichment within PSD-like condensates.

The pronounced enrichment of Tau and Fyn_SH3SH2 within the condensed phase raised the possibility that Tau-Fyn interactions influence condensate dynamics. To test this hypothesis, we monitored the mobility of Fyn_SH3SH2 within PSD 2X condensates by fluorescence recovery after photobleaching (FRAP) in the absence and presence of Tau (Fig. 2D). In condensates containing PSD-95, GKAP and Fyn_SH3SH2, Fyn_SH3SH2 displayed partial fluorescence recovery with a mobile fraction of approximately 40% (Fig. 2E,F). Strikingly, addition of Tau markedly reduced fluorescence recovery and decreased the mobile fraction of Fyn_SH3SH2 to 17% (Fig. 2E,F), indicating a pronounced dynamic arrest of Fyn_SH3SH2 within PSD-like condensates.

The sixth Tau PXXP motif, containing residues P216 and P219, represents a major interaction hotspot for Fyn and has previously been implicated in the regulation of neuronal hyperexcitability and seizure susceptibility ^24, 41, 43, 44^. To determine whether this disease-relevant interaction interface contributes to condensate dynamics, we generated a full-length human 2N4R Tau variant carrying P216A and P219A substitutions (Tau^P216A/P219A^). Mixing Tau^P216A/P219A^ with PSD 2X scaffold proteins and Fyn_SH3SH2 resulted in robust droplet formation and efficient partitioning of all components into the condensed phase (Supplementary Fig. 1), indicating that the mutations do not impair condensate incorporation.

We next examined whether disruption of the P216/P219 interaction hotspot alters the dynamic behavior of Fyn_SH3SH2 within PSD condensates. FRAP analysis revealed substantially increased fluorescence recovery of Fyn_SH3SH2 in condensates containing Tau^P216A/P219A^ compared with wild-type Tau (Fig. 2D,E). Whereas the mobile fraction of Fyn_SH3SH2 decreased from 40% to 17% upon addition of wild-type Tau, the P216A/P219A mutant restored the mobile fraction to 39% (Fig. 2F). Thus, disruption of the P216/P219 interaction hotspot almost completely reversed Tau-induced dynamic arrest of Fyn_SH3SH2.

Together, these findings demonstrate that Tau and Fyn become preferentially enriched within PSD-like condensates and reveal that a disease-relevant Tau–Fyn interaction interface is required for Tau-mediated dynamic arrest of Fyn. These results establish a mechanistic link between multivalent Tau–Fyn interactions and the regulation of condensate dynamics.

### Tau-Fyn cooperation induces dynamic arrest of PSD-95 in PSD-like condensates

Having established that Tau and Fyn co-partition into PSD condensates and that Tau–Fyn interactions promote dynamic arrest of Fyn, we next asked whether these interactions alter the dynamic behavior of the central PSD scaffold protein PSD-95. Because PSD-95 functions as a key organizational hub within the postsynaptic density ^46^, changes in its mobility provide a sensitive readout of condensate dynamics.

We first examined the effect of Tau on PSD-95 dynamics within PSD 2X condensates. In the absence of Tau and Fyn_SH3SH2, PSD-95 displayed rapid fluorescence recovery after photobleaching (FRAP) and remained highly mobile, with a mobile fraction of approximately 99% (Fig. 3A). Addition of Tau slowed PSD-95 recovery kinetics and reduced the mobile fraction to 88% (Fig. 3B-D), indicating that Tau alone partially restricts PSD-95 mobility within PSD-like condensates.

**Fig. 3.**
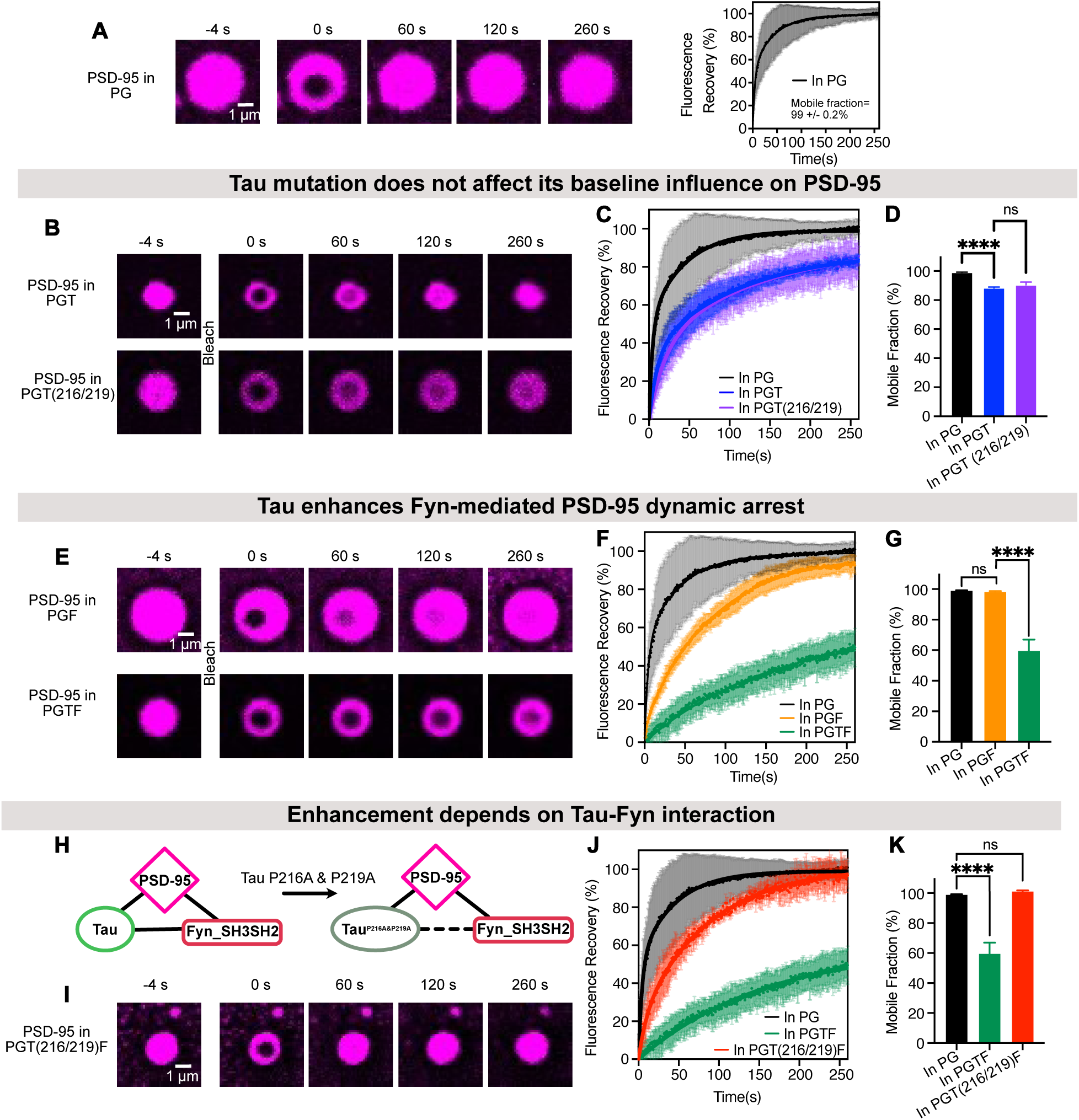
Tau-Fyn cooperation induces dynamic arrest of PSD-95 in PSD-like condensates. **A.** Representative FRAP images of 647-labelled PSD-95 in PSD-95/GKAP (“PG”) condensates (left) and quantification of 647-PSD-95 dynamics (right). Data are presented as mean values +/- SD from 4 independent experiments. The mobile fraction derived from the fitted average FRAP curve is indicated in the figure as mean value +/- SD. **B.** Representative FRAP images of 647-labelled PSD-95 in PSD-95/GKAP condensates with Tau (“PGT”; top) and with Tau^P216A/P219^ (“PGT(216/219)”; bottom). **C.** Quantification of 647-PSD-95 dynamics in PG, PGT and PGT(216/219) condensates. Data are presented as mean values +/- SD from 4 independent experiments. **D.** Mobile fractions of PSD-95 derived from fitted averaged FRAP curves in (C). Curves were averaged from 4 independent experiments. Error bars represent the SD of curve fits; one-way ANOVA: ****p ≤ 0.0001. **E.** Representative FRAP images of 647-labelled PSD-95 in PSD-95/GKAP condensates with Fyn_SH3SH2 alone (“PGF”; top) and with Tau and Fyn_SH3SH2 (“PGTF”; bottom). **F.** Quantification of 647-PSD-95 dynamics in PG, PGF and PGTF condensates. Data are presented as mean values +/- SD from n independent experiments. The number of experiments (n) for PSD-95 in PG, PGF and PGTF condensates are 4, 7, and 4, respectively. **G.** Mobile fractions of PSD-95 were derived from the fitted averaged FRAP curves in (F), which were averaged from n independent experiments. For each condition, n corresponds to the number of independent FRAP experiments indicated in (F). Error bars represent SD of curve fits; one-way ANOVA: ****p ≤ 0.0001. **H.** Schematic diagram of multivalent interactions among PSD-95, Fyn_SH3SH2 and Tau (or Tau^P216A/P219A^). Black lines indicate protein-protein interactions. The dashed line indicates weakened Tau-Fyn interaction due to P216A and P219A mutations in Tau. **I.** Representative FRAP images of 647-labelled PSD-95 in PSD-95/GKAP condensates with Tau^P216A/P219A^ and Fyn_SH3SH2 (“PGT(216/219)F”). **J.** Quantification of 647-PSD-95 dynamics in PGT, PGTF and PGT(216/219)F condensates. Data are presented as mean values +/- SD from n independent experiments. The number of experiments (n) for PSD-95 in PG, PGTF and PGT(216/219)F condensates are 4, 4, and 3, respectively. **K.** Mobile fractions of PSD-95 were derived from the fitted averaged FRAP curves in (J), which were averaged from n independent experiments. For each condition, n corresponds to the number of independent FRAP experiments indicated in (J). Error bars represent the standard deviation of curve fits; one-way ANOVA: ****p ≤ 0.0001. All FRAP curves (dots) were fitted with a bi-exponential function (solid lines).

To determine whether the proline-rich Tau-Fyn interaction hotspot contributes to this effect, we compared wild-type Tau with the Tau^P216A/P219A^ mutant. PSD-95 recovery kinetics and mobile fractions were comparable in condensates containing either wild-type Tau or Tau^P216A/P219A^ (Fig. 3B-D), indicating that the P216/P219 residues are not required for the effects of Tau alone on PSD-95 dynamics. This observation is consistent with our previous finding that Tau primarily interacts with PSD-95 through its microtubule-binding repeat region rather than through the proline-rich domain, suggesting that direct Tau-PSD-95 interactions are largely preserved in the Tau^P216A/P219A^ mutant.

We next investigated whether Fyn_SH3SH2 influences PSD-95 dynamics. Fyn_SH3SH2 modestly slowed PSD-95 recovery kinetics, reducing fluorescence recovery at 120 s from approximately 95% to 77% (Fig. 3E,F). However, the mobile fraction of PSD-95 remained largely unchanged (Fig. 3G), indicating that Fyn_SH3SH2 alone exerts only a limited effect on the equilibrium between mobile and immobile PSD-95 populations.

In contrast, simultaneous incorporation of Tau and Fyn_SH3SH2 produced a markedly stronger effect. Addition of Tau to PSD 2X/Fyn_SH3SH2 condensates substantially slowed PSD-95 recovery and reduced the mobile fraction to 59% (Fig. 3G), considerably lower than observed with Tau alone (88%; Fig. 3D). Thus, the combined presence of Tau and Fyn_SH3SH2 induced a degree of dynamic arrest that exceeded the effects of either component individually, suggesting cooperative regulation of PSD condensate dynamics by Tau and Fyn.

To directly test whether this cooperative effect depends on Tau-Fyn interactions, we disrupted the major Tau-Fyn interaction hotspot by introducing the P216A/P219A mutations into Tau and examined PSD-95 dynamics in PSD 2X/ Tau^P216A/P219A^ /Fyn_SH3SH2 condensates (Fig. 3H,I). FRAP analysis revealed substantially enhanced recovery of fluorescently labeled PSD-95 compared with condensates containing wild-type Tau and Fyn_SH3SH2 (Fig. 3J). Remarkably, the mobile fraction of PSD-95 increased from 59% in the presence of wild-type Tau and Fyn_SH3SH2 to approximately 100% in condensates containing Tau^P216A/P219A^ and Fyn_SH3SH2 (Fig. 3K). Thus, disruption of the P216/P219 interaction hotspot completely reversed the Tau–Fyn-induced dynamic arrest of PSD-95.

Together, these findings demonstrate that the pronounced dynamic arrest of PSD-95 does not arise from the independent effects of Tau or Fyn alone but instead emerges from cooperative Tau-Fyn interactions within PSD condensates. The complete rescue produced by the disease-relevant Tau^P216A/P219A^ mutant establishes the P216/P219 interaction hotspot as a critical determinant of condensate dynamics and identifies multivalent Tau-Fyn interactions as a key mechanism regulating the dynamic state of PSD-like assemblies.

### Multivalent Tau–Fyn interactions provide a structural basis for condensate dynamic arrest

Disruption of the disease-relevant P216/P219 interaction hotspot completely reversed Tau–Fyn-mediated dynamic arrest of PSD-95 and Fyn within PSD condensates. However, the ability of the Tau^P216A/P219A^ mutant to partition into condensates suggested that these mutations do not abolish Tau–Fyn interactions entirely. We therefore sought to define the molecular basis of Tau-Fyn binding and determine how disruption of P216/P219 alters this interaction network.

Because SH3 domains are the principal mediators of proline-rich motif recognition, we first characterized the interaction between full-length human 2N4R Tau and the isolated Fyn SH3 domain (Fyn_SH3) by NMR spectroscopy. Two-dimensional ^1^H-^15^N HSQC spectra of ^15^N-labeled Tau showed progressive chemical shift perturbations upon addition of 2-fold and 6-fold molar excesses of Fyn_SH3 (Fig. 4A), indicating direct interaction between Tau and the Fyn SH3 domain.

**Fig. 4.**
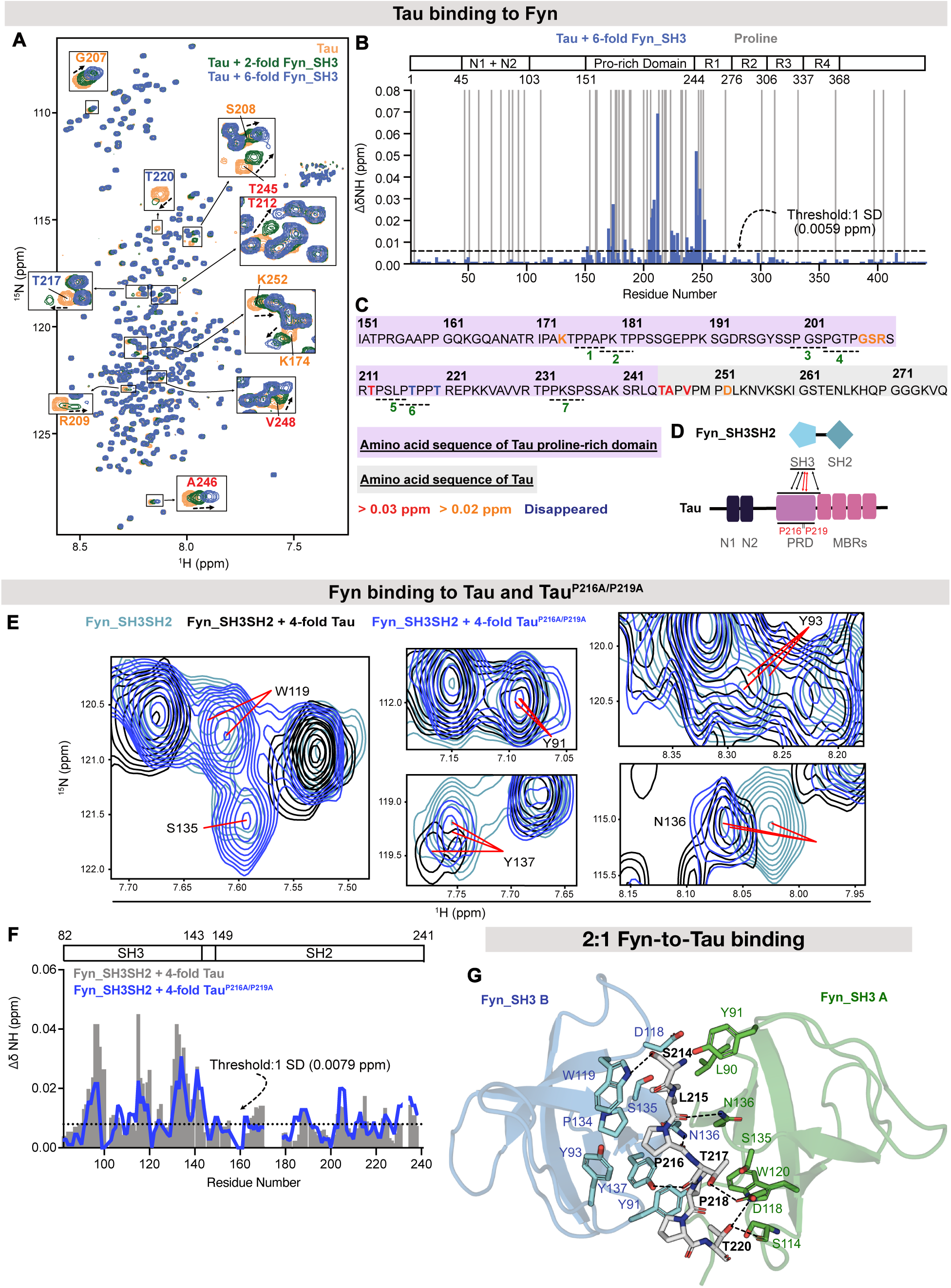
Multivalent interactions between Tau and Fyn. **A.** Superposition of ^1^H-^15^N HSQC spectra for 20 µM ^15^N-labeled 2N4R Tau alone (orange), with 2-fold (40 µM; green) and 6-fold (120 µM; blue) molar excess of Fyn_SH3. Tau residues with strong Fyn_SH3-induced chemical shift changes and peak intensity attenuation are highlighted in zoom-in views. The Tau residues for which peaks disappeared, as well as those with chemical shift changes over 0.02 ppm and 0.03 ppm, are shown in dark blue, orange, and red, respectively. **B.** NMR chemical shift perturbation analysis of 2N4R Tau upon the addition of 6-fold molar excess of Fyn_SH3. Chemical shift changes of Tau residues are shown in blue; the positions of proline residues are shown in grey. **C.** Amino acid sequence of the proline-rich domain (highlighted in purple) and R1 region (highlighted in grey) of Tau. The color code of Tau residues is the same as in (A). **D.** Schematic diagram of multivalent interactions between Fyn_SH3 and Tau. The critical residues P216 and P219 involved in Tau–Fyn interaction are highlighted in red. Red arrows indicate key Fyn_SH3–Tau interactions at the fifth and sixth PXXP motifs containing P216 and P219, whereas black arrows indicate additional interaction sites. N1 and N2 represent the N-terminal inserts of Tau; PRD represents proline-rich domain; MBRs represent the microtubule-binding repeats of Tau. **E.** Overlaid selected regions of ^1^H-^15^N HSQC spectra are shown for 50 µM ^15^N-labeled Fyn_SH3SH2 alone (cyan), with 4-fold molar excess (200 µM) of 2N4R Tau (black), and with 4-fold molar excess of Tau^P216A/P219A^ (blue). Red lines indicate the centre of Fyn_SH3SH2 cross-peaks. **F.** NMR chemical shift perturbation analysis of Fyn_SH3SH2 upon the addition of a 4-fold molar excess of Tau (grey bars) and a 4-fold molar excess of Tau^P216A/P219A^ (blue line). The black dashed line indicates the threshold for significant chemical shift changes in Fyn_SH3SH2 cross-peaks upon the addition of a 4-fold molar excess of Tau^P216A/P219A^, defined as one standard deviation (1 SD) of the shift changes. The domain organization of Fyn_SH3SH2 is shown above. **G.** Tau peptide (S214-T220) in heterotrimeric complex with two Fyn_SH3 (PDB: 9GHK). Surface and cartoon representations of Fyn_SH3 A and B are shown in green and blue, respectively. Tau peptide is shown in stick representation. Hydrogen bonds formed between Fyn_SH3 and Tau peptide are shown in black dashed lines. The figure was created using Pymol.

Mapping the chemical shift perturbations onto the Tau sequence revealed that the strongest effects were concentrated within the proline-rich region and extended into the beginning of the R1 repeat region (Fig. 4B). More detailed analysis identified particularly strong perturbations within the fifth and sixth PXXP motifs (Fig. 4C). Residues T217 and Y220 exhibited the most pronounced signal attenuations, with complete disappearance of their cross-peaks upon addition of a sixfold molar excess of Fyn_SH3 (Fig. 4A,C), indicating that the interaction is centered around the adjacent residues P216 and P219. However, additional perturbations were observed within the first PXXP motif (^176^PPAP^179^), the fourth PXXP motif (^203^PGTP^206^), and a proline-rich segment within the R1 region (^245^TAPVPMPD^252^) (Fig. 4C). These findings demonstrate that Tau engages Fyn through multiple interaction sites, with the sixth PXXP motif representing a major interaction hotspot rather than the sole binding interface (Fig. 4D).

To further characterize this interaction network, we examined binding of full-length Tau to ^15^N-labeled Fyn_SH3SH2. Addition of a fourfold molar excess of Tau induced extensive chemical shift perturbations and signal attenuations within the SH3 domain of Fyn_SH3SH2 (Fig. 4E,F). Residues forming the canonical SH3 ligand-binding groove, including Y91, Y93, W119 and Y137, displayed particularly strong perturbations (Fig. 4E,F), consistent with direct engagement of Tau. Residue S135, adjacent to the conserved ligand-binding residue P134, underwent complete signal disappearance, while N136 exhibited large chemical shift perturbations (Fig. 4E). Interestingly, peak splitting was observed for residues Y137 and N136 upon Tau binding (Fig. 4E), suggesting the coexistence of multiple binding modes. Together, these observations indicate that Tau interacts with the Fyn SH3 domain through a heterogeneous and dynamic interaction network.

We next asked how disruption of the P216/P219 hotspot alters Tau–Fyn binding. Addition of a fourfold molar excess of Tau^P216A/P219A^ still induced chemical shift perturbations and intensity changes in Fyn_SH3SH2 spectra (Fig. 4F), demonstrating that the mutant retains the ability to interact with Fyn. However, the magnitude of these perturbations was consistently reduced compared with wild-type Tau. In particular, residues W119 and S135 exhibited weaker signal attenuation, while Y91, Y93 and Y137 displayed smaller chemical shift changes in the presence of Tau^P216A/P219A^ (Fig. 4E,F). In contrast, perturbations of N136 remained largely unaffected, consistent with previous structural studies indicating that N136 primarily recognizes the backbone carbonyl group of P216 rather than its pyrrolidine side chain ^43^. These results demonstrate that the P216A/P219A mutations substantially weaken but do not abolish Tau–Fyn interactions.

The persistence of residual binding after disruption of the major interaction hotspot suggested that Tau–Fyn interactions are inherently multivalent. To obtain structural insight into this multivalent binding mode, we determined the crystal structure of the Fyn SH3 domain in complex with a Tau peptide encompassing the sixth PXXP motif (S214–T220; SLPTPPT) at 1.4 Å resolution (PDB: 9GHK). The crystal structure revealed a ternary assembly in which a single Tau peptide is sandwiched between two Fyn SH3 domains (Fig. 4G). Both Fyn_SH3 molecules engage the Tau peptide through multiple hydrogen-bonding and hydrophobic interactions (Fig. 4G and Supplementary Fig. 2), resulting in a 2:1 Fyn:Tau stoichiometry.

This architecture provides a structural framework for understanding the multivalent nature of Tau–Fyn interactions. The ability of a single Tau motif to engage multiple Fyn SH3 domains increases interaction valency and may promote the formation of highly connected interaction networks within condensates. Together with the NMR data, the crystal structure explains why disruption of P216/P219 strongly attenuates Tau–Fyn-mediated dynamic arrest while preserving residual binding: P216/P219 constitute a major interaction hotspot embedded within a broader multivalent interaction network. These findings establish a molecular and structural basis for the cooperative regulation of PSD condensate dynamics by Tau and Fyn.

## Discussion

The interaction between Tau and the Src-family kinase Fyn has emerged as a central mediator of synaptic dysfunction in Alzheimer’s disease and related tauopathies ^9, 12^. Genetic, pharmacological and biochemical studies have demonstrated that Tau is required for multiple pathological Fyn-dependent processes, including amyloid-β-induced synaptic deficits, neuronal hyperexcitability and excitotoxic signaling ^7, 8, 9, 10^. At the same time, recent work has established that the postsynaptic density (PSD) is organized through biomolecular condensation, providing a framework for understanding how large signaling networks are assembled and dynamically regulated within dendritic spines ^27, 29^. Despite these advances, it has remained unclear whether disease-associated Tau–Fyn interactions influence the dynamic organization of PSD assemblies. In the present study, we identify a condensate-based mechanism through which multivalent Tau– Fyn interactions cooperatively regulate PSD dynamics. We show that Tau and Fyn become preferentially enriched within PSD-like condensates and membrane-associated PSD clusters, that their combined presence induces a pronounced dynamic arrest of PSD condensate components, and that disruption of a disease-relevant interaction hotspot centered on residues P216 and P219 almost completely reverses these effects. Together, our findings identify PSD condensate dynamics as a previously unrecognized target of pathological Tau–Fyn signaling and establish a mechanistic framework linking a disease-relevant Tau–Fyn interaction hotspot to the regulation of postsynaptic molecular organization.

Our findings extend current models of Tau–Fyn signaling in several important ways. Tau has traditionally been viewed primarily as a trafficking and scaffolding factor that recruits or stabilizes Fyn at postsynaptic sites, thereby facilitating phosphorylation-dependent modulation of NMDAR signaling (Fig. 5) ^9^. While this model is supported by substantial experimental evidence ^9^, it does not readily explain how Tau influences the organization and dynamics of the highly interconnected signaling networks that characterize the PSD. Our data suggest that Tau may contribute to postsynaptic dysfunction not only by influencing the localization of Fyn but also by modulating the material properties of PSD assemblies themselves (Fig. 5). We find that Tau and Fyn are preferentially enriched within PSD condensates relative to core scaffold proteins and that their combined presence markedly reduces the mobility of both Fyn and PSD-95. These observations suggest that pathological Tau–Fyn signaling may influence synaptic function by altering the dynamic exchange of proteins within postsynaptic signaling assemblies.

**Fig. 5.**
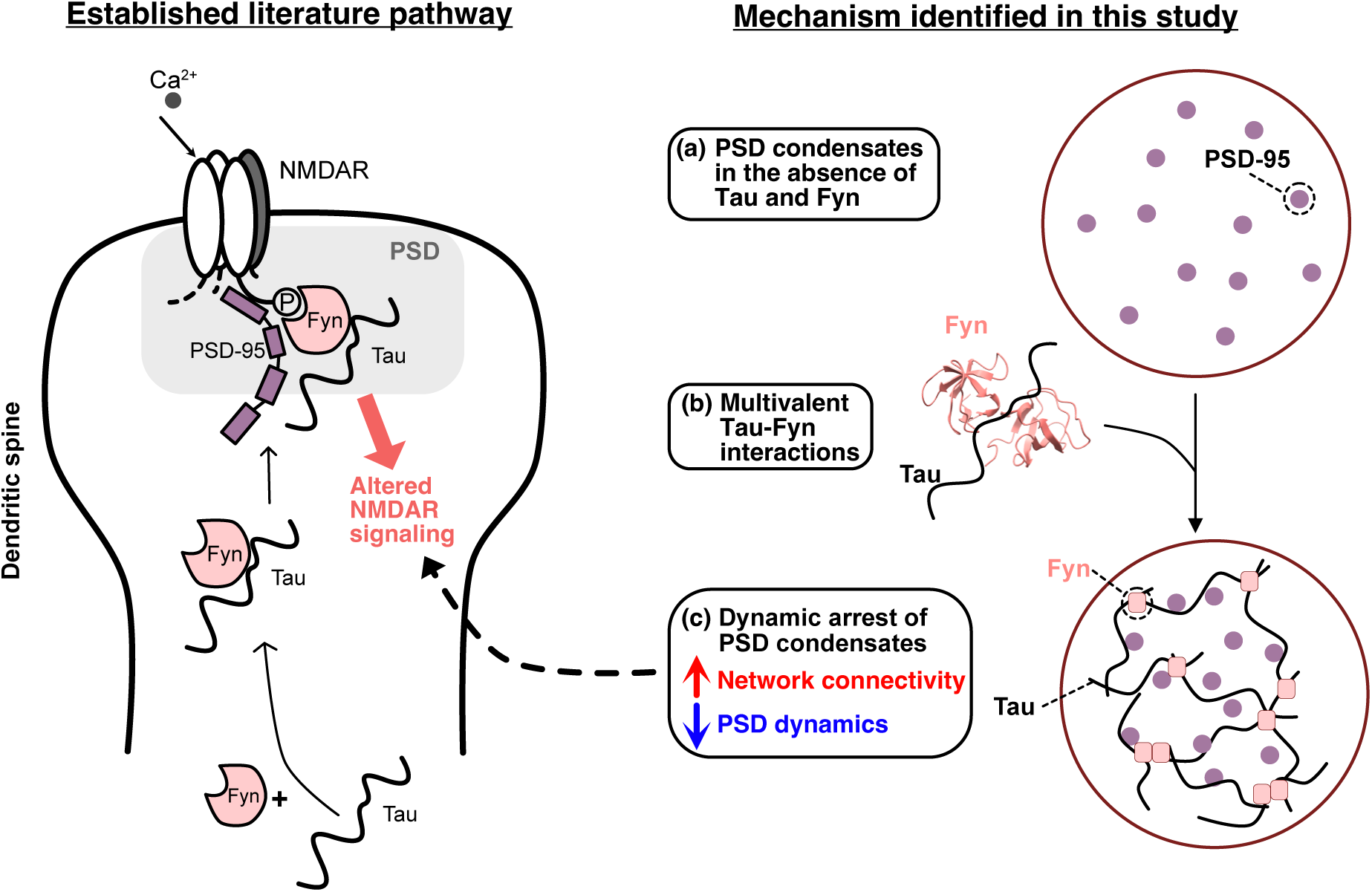
Proposed model for cooperative regulation of PSD organization by Tau and Fyn. Previous studies (left) have shown that Tau recruits the tyrosine kinase Fyn to the postsynaptic density (PSD), where it phosphorylates Y1472 on the NMDA receptor subunit NR2B, leading to altered NMDAR signaling and excitotoxicity ^9^. In the present study (right), we identify a distinct mechanism whereby Tau and Fyn have been enriched in the PSD condensates and multivalent Tau–Fyn interactions increase network connectivity within the PSD condensate. This enhanced connectivity is associated with reduced PSD-95 mobility and dynamic arrest of PSD condensates. These findings suggest that Tau–Fyn interactions can alter the dynamic properties of PSD condensates and thereby influence the molecular environment of the postsynaptic density. The dashed arrow denotes a potential connection between condensate arrest and altered NMDAR signaling.

A notable aspect of our findings is that these effects are mediated by a kinase-deficient Fyn_SH3SH2 construct lacking the catalytic domain of Fyn. Thus, the observed regulation of condensate dynamics does not require Fyn kinase activity and instead arises from the adaptor functions of Fyn mediated by its SH3 and SH2 domains. These observations reveal a previously underappreciated kinase-independent role of Fyn as a structural regulator of postsynaptic organization. This function complements, rather than replaces, the well-established kinase-dependent role of Fyn in NMDAR signaling and suggests that Fyn contributes to synaptic regulation through both enzymatic and architectural mechanisms.

More broadly, our findings extend emerging concepts in the PSD condensate field. Previous studies have primarily focused on how core scaffold proteins such as PSD-95, GKAP, Shank and Homer assemble PSD condensates and organize receptor-associated signaling complexes ^29^. In contrast, considerably less is known about how disease-associated signaling proteins remodel the dynamic properties of these assemblies once formed. Our results demonstrate that a neurodegeneration-associated interaction network can profoundly alter PSD condensate behavior without being required for condensate formation itself. Thus, Tau and Fyn function not merely as clients of PSD condensates but as active regulators of condensate dynamics. This observation supports the broader concept that pathological signaling proteins may contribute to disease by remodeling the material properties of biomolecular condensates.

A particularly notable finding is that disruption of the sixth Tau PXXP motif almost completely restores condensate dynamics despite only partially weakening Tau–Fyn binding. Previous studies demonstrated that mutation of residues P216 and P219 rescues neuronal hyperexcitability, seizure susceptibility and sleep abnormalities in vivo, identifying this motif as a critical determinant of Tau-dependent pathology ^24^. Our data provide a potential molecular explanation for these observations. NMR spectroscopy demonstrates that Tau engages Fyn through a distributed interaction network involving multiple PXXP motifs and additional proline-rich segments, whereas P216 and P219 constitute a dominant interaction hotspot within this network. Consequently, mutation of P216 and P219 attenuates but does not abolish Tau–Fyn binding. The near-complete restoration of condensate dynamics by the P216A/P219A mutations therefore suggests that the ability of Tau–Fyn interactions to generate highly connected interaction networks may be more important for condensate regulation than binding affinity alone. This distinction may help explain why selective disruption of a single interaction hotspot produces disproportionately large functional effects in cellular and animal models ^24^.

An important feature of our data is that the pronounced arrest of PSD-95 emerges only when Tau and Fyn are present together. Tau alone partially reduces PSD-95 mobility, whereas Fyn alone exerts comparatively modest effects. In contrast, the combination of Tau and Fyn produces substantially stronger dynamic arrest than either protein individually, and this effect is completely reversed by disruption of the P216/P219 interaction hotspot. These findings indicate that dynamic arrest is an emergent property of cooperative Tau–Fyn interactions rather than the additive consequence of independent Tau and Fyn activities. Such emergent behavior is a hallmark of multivalent interaction networks and is consistent with the highly interconnected architecture of biomolecular condensates.

One mechanistic interpretation of these observations is that Tau–Fyn complexes function as multivalent crosslinkers within PSD assemblies. In this model, the distributed interaction network formed between Tau and Fyn increases the effective connectivity of the PSD interaction network and reduces molecular exchange within the condensate (Fig. 5). The resulting increase in network connectivity would be expected to promote transitions toward more dynamically arrested or gel-like states (Fig. 5). Consistent with this interpretation, disruption of the dominant P216/P219 interaction hotspot completely restores PSD-95 mobility despite only partially reducing Tau–Fyn binding. These findings suggest that condensate regulation depends primarily on interaction valency and network connectivity rather than on binding affinity alone.

The structural data provide additional mechanistic insight into this phenomenon. The observation that a single Tau peptide can simultaneously engage two Fyn SH3 domains supports a model in which Tau increases interaction valency within PSD assemblies. Such multivalent interactions are a defining feature of biomolecular condensates and frequently drive transitions between more dynamic and more arrested material states. Although the precise stoichiometries and geometries present within condensates are unlikely to be fully captured by a crystallographic assembly, the structure reveals an interaction architecture that is consistent with the multivalent binding behavior observed by NMR spectroscopy and with the strong effects of Tau–Fyn interactions on condensate dynamics. Rather than representing a definitive structural model of the condensate state, the crystal structure provides a plausible molecular mechanism through which Tau may enhance network connectivity within PSD assemblies.

The observed changes in PSD-95 dynamics may also have broader implications for postsynaptic signaling. PSD-95 serves as a central scaffold linking NMDARs and other receptors to downstream signaling pathways, and dynamic exchange of PSD proteins is thought to regulate receptor accessibility, signaling complex assembly and activity-dependent remodeling of synaptic connections. Previous work demonstrated that alterations in PSD condensate dynamics influence the organization of receptor-associated signaling complexes, and we previously showed that Tau alone can slow the dynamics of PSD proteins through direct interactions with PSD-95 ^28^. The present study extends these findings by demonstrating that Tau and Fyn cooperatively induce substantially stronger dynamic arrest than either protein alone. Although our experiments do not directly measure NMDAR activity, receptor trafficking or synaptic signaling, they identify a mechanism through which pathological Tau–Fyn interactions could alter the molecular environment in which NMDAR signaling occurs. One possibility is that prolonged residence times of scaffold and signaling proteins within PSD assemblies stabilize receptor-associated signaling complexes and thereby influence their responsiveness to synaptic activity. Future studies will be required to determine how the condensate dynamics described here influence receptor phosphorylation, receptor mobility and downstream signaling events in living neurons.

The PSD contains a dense network of SH3 domain–proline-rich domain interactions involving numerous scaffold and signaling proteins. For example, Tau interacts with BIN1, a major genetic risk factor for Alzheimer’s disease implicated in receptor trafficking, while Shank proteins contain extensive proline-rich regions that contribute to PSD organization and are frequently mutated in neurodevelopmental disorders ^47, 48, 49, 50, 51^. The multivalent Tau–Fyn interaction network described here is therefore likely embedded within a broader interaction landscape. An important direction for future work will be to determine how Tau–Fyn complexes interact with competing or cooperative SH3-mediated signaling networks and whether disease-associated alterations in these networks further modulate PSD condensate dynamics.

An additional layer of regulation may arise from post-translational modifications. Tau phosphorylation within the proline-rich region has been shown to modulate Tau–Fyn interactions, and Fyn itself phosphorylates Tau at multiple sites, including Y18 ^43, 52, 53, 54^. Such modifications could potentially alter interaction valency by creating additional SH2-mediated binding interfaces ^55^ or by changing the affinity of existing SH3-dependent interactions. Although these possibilities were not examined in the present study, they suggest that Tau–Fyn-mediated condensate regulation may itself be dynamically controlled by disease-associated signaling pathways. The potential existence of such feedback mechanisms represents an intriguing area for future investigation.

Several limitations should be considered when interpreting our findings. First, the experiments were performed in reconstituted systems designed to capture key features of PSD organization while reducing the complexity of native synapses. Such systems enable quantitative dissection of molecular mechanisms but do not fully reproduce the biochemical and structural complexity of neuronal postsynaptic densities. Second, our study focuses primarily on the adaptor functions of Fyn mediated by its SH3 and SH2 domains and does not address potential contributions of kinase activity to condensate regulation. Third, although the strong concordance between our mutational, biophysical and structural data supports the physiological relevance of the identified interaction hotspot, direct validation in neurons will be necessary to establish how Tau–Fyn-mediated condensate regulation contributes to synaptic function and dysfunction in vivo.

Despite these limitations, the convergence of condensate reconstitution, quantitative partitioning analysis, mutagenesis, NMR spectroscopy and crystallography provides a coherent mechanistic model for how Tau and Fyn cooperate within postsynaptic assemblies. More broadly, our findings support the emerging view that neurodegeneration-associated proteins can influence cellular function not only through canonical signaling pathways but also through regulation of biomolecular condensate dynamics. We propose that multivalent Tau–Fyn interactions act as a molecular rheostat that modulates the dynamic state of PSD assemblies. More generally, these findings suggest that pathological remodeling of condensate dynamics may represent a previously unrecognized mechanism through which Tau perturbs postsynaptic signaling in Alzheimer’s disease and related tauopathies.

## Materials and Methods

### Protein expression

For PSD proteins used in this study, the previously described design of the protein constructs was followed ^29^: full-length PSD-95; PSD-95_SH3-GK containing the SH3 and GK domains; GKAP_GBR_PBM containing three GK-binding repeats (GBR) and the PDZ-binding motif (PBM); Shank_M1718E_PDZ_HBS_CBS_SAM with M1718E mutation and containing the PDZ, Homer-binding sequence (HBS), cortactin-binding sequence (CBS) and sterile alpha motif (SAM); full-length Homer (residues M1-P361); His_6_-tagged GCN4-NR2B. For Fyn constructs, the design followed ^56^: Fyn_SH3 containing the SH3 domain; Fyn_SH3SH2 consisting of the SH2 and SH3 domains. The three cysteine residues were mutated into serine in the Fyn_SH3SH2 construct to avoid non-specific disulfide bond formation. Full-length 2N4R Tau constructs without any mutations and with P216A and P219A were used.

The DNA of each PSD or Fyn protein construct was ordered from Invitrogen GeneArt Gene Synthesis Services (ThermoFisher Scientific) and cloned into a modified pET 28a vector (with a Z2 solubility tag and N-terminal His_6_-Tev tag). For human 2N4R Tau (htau40; UniProt: P10636-8), its gene was inserted into the pNG2 vector (a derivative of pET-3a, Novagen). In all cases, plasmids with genes of interest were transformed into the *E. coli* BL21 (DE3) cells. Recombinant proteins were expressed in Escherichia coli BL21 (DE3) cells (Novagen).

For unlabeled proteins, cells were grown in LB medium supplemented with kanamycin, and protein expression was induced by 0.5 mM IPTG (1 mM for Shank and Homer) when OD600 reached 0.8. After protein expression at 16°C overnight (or 30 °C for 3 hrs for PSD-95_SH3-GK; 37 °C for 3 h for His-NR2B; and 25 °C overnight for Shank), cells were harvested by centrifugation with 7,000 g at 4 °C for 30 min.

For expression of uniformly ^15^N-labeled Fyn_SH3SH2 and ^15^N-labeled 2N4R human Tau, the protein was expressed until OD600 of 0.6 to 0.8 was reached. The cells from 1 L LB were pelleted by centrifugation with 5000 g at 20 °C for 30 min, followed by washing and pelleting in 500 mL 1 x M9 salt solution (3 g KH_2_PO_4_, 0.5 g NaCl, 6.78 g Na_2_HPO_4_). The cell pellet was resuspended in 250 mL of M9 minimal medium supplemented with 2g ^15^NH_4_Cl and 0.5 g ISOGRO-15N as the only nitrogen source. After 1 h of incubation at 37 °C, protein expression was induced with 0.5 mM IPTG overnight at 30 °C for ^15^N-labeled Fyn_SH3SH2 or 1 mM IPTG overnight at 37 °C for ^15^N labeled htau40, followed by centrifuging with 7000 g at 4 °C for 30 min to harvest cells.

### Protein purification

For protein purification of PSD and Fyn protein constructs, cell pellets were resuspended in lysis buffer consisting of 50 mM Tris, pH 8.0, 300 mM NaCl, 10 mM imidazole, 2 mM β-mercaptoethanol (6 mM β-mercaptoethanol for PSD-95), 0.5 mM PMSF, 1 mg/mL lysozyme, 5 µg/mL DNase, 1 mM MgCl_2_, and cOmplete EDTA-free protease inhibitor cocktail. The cells were disrupted by sonication on ice. After sonication, cellular debris were removed by centrifugation at 48,254 x g and 4 °C for 40 min. Target proteins present in the supernatant were purified by immobilized metal affinity chromatography using a Ni^2+^ affinity column (GE Healthcare or Qiagen) equilibrated with 50 mM Tris, pH 8.0, 300 mM NaCl, 10 mM imidazole, and 2 mM β-mercaptoethanol (or 6 mM β-mercaptoethanol for PSD-95). Non-specifically bound proteins were removed by washing the column after loading with this buffer until a stable baseline was reached again. Specifically bound protein was finally eluted with 50 mM Tris, pH 8.0, 300 mM NaCl, 300 mM imidazole, and 2 mM β-mercaptoethanol (or 6 mM β-mercaptoethanol for PSD-95) as elution buffer. The eluted proteins were dialyzed overnight at 4 °C against TEV cleavage buffer (50 mM Tris, pH 8.0, 150 mM NaCl, 0.1 mM EDTA, 0.5 mM PMSF, 2 mM β-mercaptoethanol and 6 mM β-mercaptoethanol for PSD-95) and cleaved by TEV the next day. After TEV cleavage, non-cleaved proteins and residual His_6_-tags were removed from the sample using a Ni^2+^ affinity column and collecting the flow through. The target proteins were concentrated and further purified by size-exclusion chromatography (Superdex 75 or Superdex 200 columns; GE Healthcare) using a buffer containing 50 mM Tris, pH 8.0, 300 mM NaCl, 1 mM EDTA, and 3 mM DTT. For the purification of PSD-95, PSD-95_SH3-GK, Homer and His-NR2B, the proteins were further purified by ion exchange chromatography to remove unwanted contaminants. In addition, His-NR2B purification skipped the TEV cleavage and the second Ni^2+^ affinity chromatography (after TEV cleavage) steps to keep the N-terminal His_6_-tag.

For the preparation of 2N4R human Tau and 2N4R human Tau mutant P216A/P219A-Tau, bacterial cells were harvested by centrifugation and the pellets were resuspended in a 20 mM MES buffer at pH 6.8, including 1 mM EGTA, 0.5 mM MgCl_2_, 1 mM PMSF, 1 mg/mL lysozyme, 10 µg/mL DNase I, 5 mM DTT and cOmplete EDTA-free protease inhibitor cocktail. Cells were disrupted by French Press on ice. The resulting lysates were supplemented with NaCl to a final concentration of 500 mM and boiled at 98 °C for 20 min. Lysates were cooled down on ice and ultracentrifuged at 127,000 x g and 4 °C for 40 min. DNA of the supernatant was precipitated by adding 20 mg/mL of streptomycin sulfate and incubated at 4 °C for 15 min with rotation. After centrifugation at 15,000 x g and 4 °C for 30 min the pellet was discarded and the resulting supernatant was incubated with 361 mg/mL (NH_4_)_2_SO_4_ at 4 °C for 15 minutes to precipitate Tau. Then, precipitated proteins were pelleted by repeating the previous centrifugation step. The pellets containing the protein were resuspended in dialysis buffer (20 mM MES buffer, pH 6.8, 1 mM EDTA, 0.1 mM PMSF, and 2 mM DTT) and dialyzed against the same buffer at 4 °C overnight to remove salts. Filtered dialysate was purified by cation exchange chromatography using a MonoS 10/100 column (Cytiva) with a linear gradient from 0 % to 60 % of elution buffer (binding buffer: 20 mM MES, pH 6.8, 50 mM NaCl, 1 mM EDTA, 2 mM DTT, and 0.1 mM PMSF; elution buffer with the same composition as binding buffer, except with 1 M NaCl). After identifying the fractions containing Tau with SDS-PAGE, these were pooled together and concentrated by ultrafiltration (5 kDa MWCO Vivaspin concentrator, Sartorius). Unlabeled 2N4R human Tau and the unlabeled 2N4R human Tau P216A/P219A mutant were further purified by reverse-phase chromatography on a preparative C4 column (Vydac 214TP, 5 µm, 8 × 250 mm) using an HPLC system. Protein purity was confirmed by electrospray ionization mass spectrometry. The purified proteins were lyophilized and redissolved in the appropriate buffer before use in subsequent experiments.

Following purification on the Mono S 10/100 column, ^15^N-labeled 2N4R human Tau was further purified by size-exclusion chromatography using a column equilibrated with PBS supplemented with 500 mM NaCl, 1 mM DTT, and 0.1 mM PMSF. Purified proteins were dialyzed against the respective buffers and protein concentrations were determined using the Pierce™ BCA Protein Assay Kit (Thermo Scientific) before the proteins were used in subsequent experiments.

All purified proteins were flash frozen and stored at −80 °C until further use.

### Phase separation assays for fluid PSD condensates

For phase separation assays, proteins were labeled with Alexa Fluor^TM^ 488, Alexa Fluor^TM^ 594, and Alexa Fluor^TM^ 647 microscale protein labeling kits (ThermoFisher Scientific, Invitrogen).

For fluorescent microscopy, small amounts of fluorescently labeled PSD-95, Tau, Tau^P216A&P219A^, Fyn or Fyn_SH3SH2 (∼ 0.3 µL) were pre-mixed with unlabeled PSD-95, Tau, Tau^P216A&P219A^, Fyn or Fyn_SH3SH2 protein stocks, respectively. Then, PSD scaffold proteins, Tau (or Tau^P216A&P219A^), Fyn (or Fyn_SH3SH2) were mixed at equal molar ratios and diluted in the phase separation buffer (50 mM Tris, pH 7.8, 100 mM NaCl, 1 mM EDTA, and 5 mM DTT) to reach the indicated protein concentrations. No condensate-inducing crowding agent was used in the condensate experiments. For each sample, 5 µL of the reaction was added onto a microscope slide with an 18 mm coverslip. DIC and fluorescent imaging were performed on a Leica DM6B microscope, and a 63x water objective was used to observe and acquire images at room temperature. Micrographs were analyzed using Fiji (NIH).

### Fluorescence recovery after photobleaching (FRAP) on droplets

The dynamic nature of phase-separated PSD and PSD/NR2B droplets was analyzed using FRAP. Phase separation of PSD proteins was induced as described previously.

FRAP was recorded on a Leica TCS SP8 confocal microscope using a 63x oil objective at room temperature. The circular region of interest (ROI) with a diameter of 1 µm was selected in the middle of each droplet. After recording 3 frames (with a frame rate of 523 ms), the fluorescence of Alexa 488-labeled Tau, Alexa 594-labeled Fyn_SH3SH2 and Alexa 647-labeled PSD-95 in ROI was photobleached by a 488-argon, a DPSS 561 laser beam and a HeNe 633 laser beam at 50 % laser power for 5 frames, respectively. After photobleaching, 500 frames were recorded to capture the fluorescence recovery. The images were analyzed using Fiji (NIH). Each FRAP curve was calculated according to:

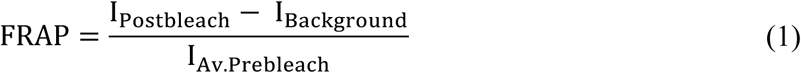

where I_Postbleach_ indicates the measured fluorescence intensity of ROI after bleaching, I_Av.Prebleach_ indicates the averaged intensity of ROI before bleaching. Both intensities were corrected by background subtraction.

The calculated results were further corrected by multiplying the acquisition bleaching correction factor (ABCF), which was calculated according to:

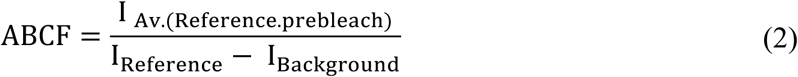

where I_Reference_ indicates the fluorescence intensity of the whole droplet area, I_Av.(Reference.prebleach)_ indicates the averaged I_Reference_ before bleaching. The values were corrected by background subtraction. Intensity differences between frames before (I_Prebleach_) and just after bleaching (I_Postbleach_ at time 0; called as I_i_) were normalized to 100 % according to:

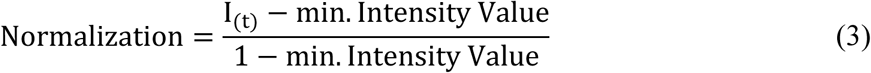

Subsequently, FRAP curves were averaged and fitted with a bi-exponential function using GraphPad Prism:

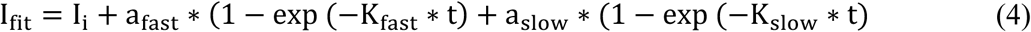

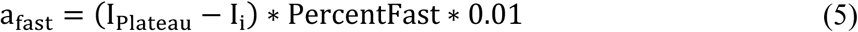

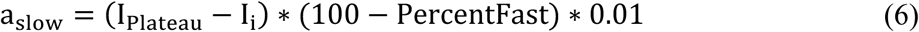

where I_i_ were constrained to a constant value of 0. I_plateau_ indicates the fluorescence intensity at infinite times and was constrained to be less than 100. K_fast_ and K_slow_ indicate the rate constants of the faster and slower component, respectively.

Mobile fractions were derived from curve fits, based on the following equation:

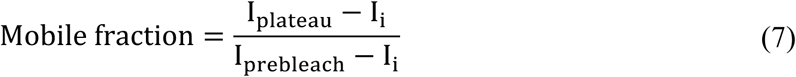

Welch’s t-test was performed to compare the mobile fractions of Fyn_SH3SH2 without and with Tau; of Tau without and with Fyn_SH3SH2 (****p ≤ 0.0001). For other experiments (as indicated), one-way ANOVA analyses were performed (**p ≤ 0.0021, ***p ≤ 0.0002, ****p ≤ 0.0001).

### Membrane-anchored PSD clustering

Phospholipids including POPC, 4 % Ni^2+^-NTA-DGS, 0.1 % PEG5000PE, and 0.1 % rhodamine-PE were mixed in chloroform in glass bottles and dried under vacuum for 2 h. 500 µL to 1 mL buffer (50 mM Tris-HCl, pH 7.8, 100 mM NaCl) was added to the dried lipid film to reach a final lipid concentration of 1 mM, and resuspended with 180 rpm shaking at 37 °C for 1 hr. The lipid mixture was then transferred to an Eppendorf tube and went through freeze-thaw for 15 runs until the lipid mixture became clear to form small unilamellar vesicles (SUVs). The SUVs were further clarified by centrifugation at 17,000 x g for 30 min and stored at 4 °C for one week.

For preparation of supported lipid bilayers, glass-bottomed 96-well plates (Greiner, Product No. 655891) were pre-cleaned with 2 % Hellmanex II overnight and 6 M NaOH for 30 min twice at room temperature. Before adding SUVs, wells were equilibrated with buffer (50 mM Tris-HCl, pH 7.8, 100 mM NaCl) for 5 min, and left 60 µL buffer in the well. 20 µL 1 mM SUVs were added and incubated for 20 min. 20 µL 5 M NaCl was then added for another 20 min for the SUVs to further collapse onto the glass bottom. Excess SUVs were washed away by pipetting in and out buffer (50 mM Tris-HCl, pH 7.8, 100 mM NaCl) eight times. The quality of supported lipid bilayers was controlled by FRAP under the confocal microscope.

Supported lipid bilayers with a certain amount of Ni^2+^-NTA-DGS lipid were blocked with 1 mg/mL BSA for 30 min. 500 nM NR2B receptors were added and incubated for 30 min. Excessive receptors were washed away by pipetting in and out buffer (50 mM Tris-HCl, pH 7.8, 100 mM NaCl, 1 mM TCEP) eight times. 1 µM of each PSD 4X scaffold protein was added with and without a certain amount of Tau for 15 min to form membrane-associated PSD clusters. Three images for each well were taken randomly under the confocal microscope. Time lapses were taken with a frame rate of 30 s for 15 min under the confocal microscope (Abberior Instruments, Göttingen, Germany).

For the partition assay with Tau/Fyn_SH3SH2, 1 µM of each PSD 4X scaffold protein with Tau/Fyn_SH3SH2 were added to supported lipid bilayers functionalized with NR2B receptors for 15 min. Three images for each well were taken randomly under the confocal microscope.

### NMR spectroscopy

To investigate the binding sites on Fyn_SH3SH2 by Tau or P216A/P219A-Tau, 2D ^1^H-^15^N HSQC experiments were recorded at 298 K on a 700 MHz spectrometer in the buffer containing 50 mM sodium phosphate, pH 6.5, 100 mM Na_2_SO_4_, 1 mM DTT and 10 % D_2_O. The spectrum was recorded with 64 scans. We set 2048 and 64 increments in the ^1^H and ^15^N dimensions, respectively. 50 µM ^15^N-labeled Fyn_SH3SH2 was titrated with 2-fold (100 µM) and 4-fold (200 µM) full-length 2N4R Tau.

In order to explore the interaction motifs of Tau binding to Fyn_SH3, 2D ^1^H-^15^N HSQC experiments were recorded at 278 K on 800 MHz and 1.2 GHz NMR spectrometers in the buffer containing 50 mM sodium phosphate, pH 6.5, 100 mM Na_2_SO_4_, 1 mM DTT and 10 % D_2_O. 20 µM ^15^N-labeled 2N4R Tau were titrated with 2-fold (40 µM) and 6-fold (120 µM) Fyn_SH3. The spectrum was recorded with 48 scans. we set 2048 and 256 increments in the 1H and 15N dimensions, respectively.

All spectra were processed with TopSpin 4.0.6 and analyzed using Sparky ^57^. Assignments of ^1^H-^15^N cross peaks of Tau and Fyn_SH3SH2 were previously determined ^58, 59^.

Chemical shift perturbation (CSP) was calculated according to:

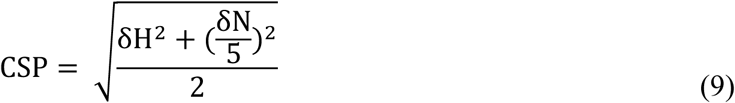

where δH and δN represent the chemical shift changes of proton and nitrogen nuclei, respectively. The CSP plot for ^15^N-Fyn_SH3SH2/Tau was further smoothed by applying a sliding average of three neighbouring residues. Because proline residues are not detectable by ^1^H-^15^N HSQC experiments and the CSP of every proline residue was set to 0, the CSP plot for ^15^N-Tau/Fyn_SH3 was not further smoothed to avoid the underestimation of chemical shift changes in proline-rich region of Tau, which is important for Tau/Fyn interactions.

### X-ray crystallography

For crystallization the concentration of the Fyn SH3 domain in 10 mM Tris/HCl, pH 8.0, was adjusted to 2 mM (14 mg/ml) and Tau (211 - 220) peptide or Tau (211 - 223) peptide was added in 2-fold or 4-fold molar excess. Crystals were obtained at 20 °C for the setups with the 2-fold and 4-fold excess of Tau (211 - 220) peptide by the vapor-diffusion technique with sitting drops composed of 100 nl of protein solution and 100 nl of well solution. The well solutions contained 0.2 M NaCl, 0.4 M NaH_2_PO_4_, 1.48 - 1.6 M K_2_HPO_4_ and 0.1 M imidazole, pH 7.0 - 8.0. Crystals were cryoprotected by transferring them for one minute to well solution supplemented with 32 % glycerol and flash-cooled by plunging them into liquid nitrogen.

Diffraction data were collected on beamline X10SA, SLS, Switzerland (Supplementary Table 1). Diffraction images were integrated and scaled using XDS ^60^. The Matthews coefficient (VM) was calculated to be 1.94 Å3 Da^-1^, corresponding to two monomers per asymmetric unit with an estimated solvent content of 36.56% ^61^. The structure of the Fyn SH3-Tau (211-220) complex was solved by molecular replacement with Phaser ^62^ using the crystal structure of the human Fyn SH3 domain ^63^ (PDB code: 3UA6). A single solution was found with two SH3 domains in complex with the Tau peptide. Refinement was performed with Refmac^64^ alternating with model building in Coot ^65^. The refinement statistics are presented in Supplementary Table 2. The atomic coordinates and structure factors have been deposited in the Protein Data Bank under accession code 9GHK.

### Determination of protein concentrations in dilute phase using HPLC

All four proteins were mixed together to a final concentration of 25 μM each in the phase separation buffer (50 mM Tris, pH 7.8, 100 mM NaCl, 1 mM EDTA, and 5 mM DTT), and incubated at 20 °C for 20 min. Samples were then centrifuged at 11,000 rpm for 5 minutes. 25 μL of the dilute phase was carefully removed without disturbing the dense phase and injected onto a reverse-phase C4 column (ReproSil Gold 300, Dr. Maisch). HPLC measurements were performed using a Jasco system equipped with an autosampler (AS-2055), pump (PU-2080), and multiwavelength detector (MD-2010 Plus), monitoring absorbance at 220 nm. The column was equilibrated in H₂O containing 0.1% trifluoroacetic acid (TFA) and 5% acetonitrile. Proteins were eluted using a linear gradient from 5% to 95% acetonitrile.

Calibration curves were generated for each protein of known concentrations. The relationship between peak area and protein concentration is described by:

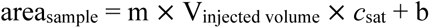

where area_sample_ is the integrated peak area, V_injected volume_ is the injected volume, and m and b are the slope and intercept of the calibration curve, respectively.

Peak areas were determined using the instrument’s built-in software, and concentrations were calculated based on the corresponding calibration curves. For further details on the analytical HPLC method for biomolecule quantification and concentration determination, see ref ^66^. At least three independent replicates were measured for each sample and averaged.

## Acknowledgement

We thank Kerstin Overkamp for protein synthesis work, Adriana Savastano and Sára Joana Varga for assistance with FRAP assays, and the light microscopy facility at the Max Planck Institute for Multidisciplinary Sciences for microscope access. M.Z. was supported by the European Research Council (ERC) under the EU Horizon 2020 research and innovation programme (grant agreement No. 787679). M.Z. acknowledges access to the 1.2 GHz spectrometer through the DFG Major Instrumentation Grant INST 1525/26-1 FUGG (project number 600373). A.H. was supported by the Deutsche Forschungsgemeinschaft (DFG), project number 402723784-SPP2191.

## Author contributions

Z.S. produced protein, performed biochemical/biophysical experiments, phase separation assays, and NMR spectroscopy. A.B. measured protein concentrations in the dilute phase using analytical HPLC. D.S. performed membrane-anchored PSD/NR2B clustering assays. M.Z. supervised biochemical/biophysical experiments, phase separation assays, and NMR spectroscopy. M.-S.C-O. prepared recombinant protein. A.H. supervised membrane-anchored PSD/NR2B clustering assays. S.B. performed X-ray crystallography experiments. All authors contributed to manuscript preparation. M.Z. designed the project.

## Competing interests

The authors declare no competing interests.

## Data availability

All data needed to evaluate the conclusions in the paper are present in the paper and/or the Supplementary Materials.

## Notes

### Competing Interest Statement

The authors have declared no competing interest.

