## Supplemental figures and tables for "Multivalent Tau-Fyn interactions cooperatively arrest postsynaptic density condensate dynamics"

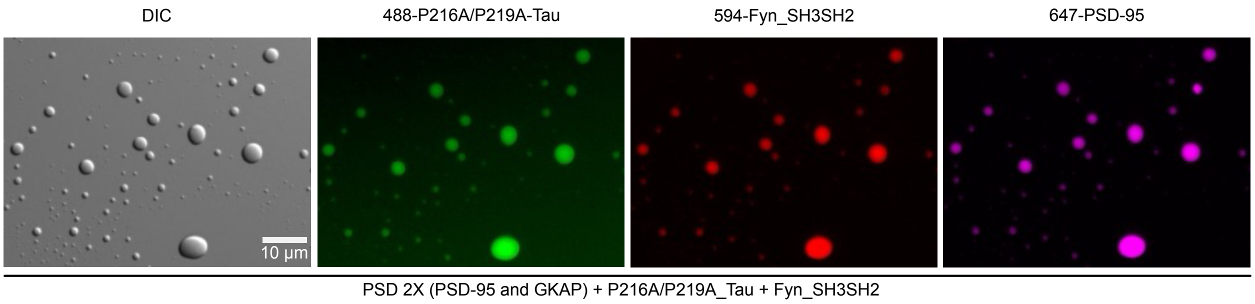


**Supplementary Fig. 1 | Tau$^{P216A/P219A}$ participates into the PSD 2X/ Tau$^{P216A/P219A}$/Fyn_SH3SH2 condensates.** DIC and fluorescence microscopy of PSD condensates containing PSD-95, GKAP, Tau$^{P216A/P219A}$ and Fyn_SH3SH2. Tau$^{P216A/P219A}$, Fyn_SH3SH2, and PSD-95 were labeled with Alexa 488, Alexa 594 and Alexa 647, respectively. 25 μM of each protein was used.

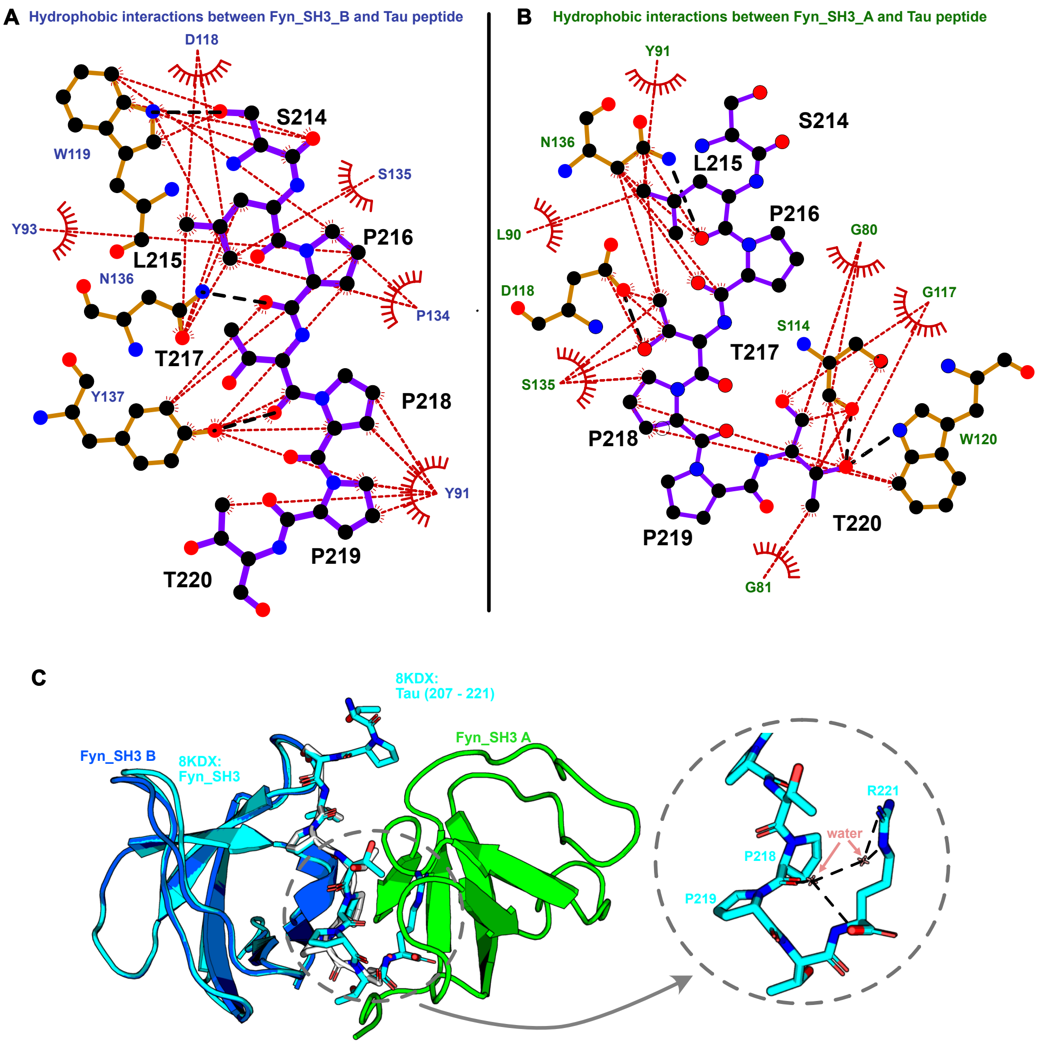


**Supplementary Fig. 2 | Atomic-resolution view of Tau-Fyn interaction. A,B.** 2D representation of the protein-peptide interaction between Tau peptide and Fyn_SH3_A (C) and Fyn_SH3_B (D). Purple, Tau peptide; brown, Fyn_SH3 residues making hydrogen bond and hydrophobic interactions; red rays, residues of Tau and Fyn_SH3 making hydrophobic interactions; red dashed lines, hydrophobic interactions; black dashed lines, hydrogen bonds; blue, red and black spheres represent nitrogen, oxygen and carbon atoms, respectively. The figures were created using LigPlot+. **C.** Cartoon representations of Fyn_SH3 domain B, Fyn_SH3 domain A, and the published binary complex (PDB: 8KDX) are shown in blue, green, and cyan, respectively. Our ternary structure (PDB: 9GHK) reveals a 2:1 protein–peptide complex, in contrast to the previously reported 1:1 stoichiometry between Fyn_SH3 and a longer Tau peptide (G207–R221)[7] . The observed difference may be influenced by the distinct peptide boundaries used in the two studies. In our shorter peptide (S214-T220), the C-terminus remains extended, exposing additional interfaces that enable recruitment of a second protein chain. Stick representations of the longer peptide (residues 207–221) from the published 8KDX binary structure and the shorter peptide (residues 214–220) are shown in cyan and white, respectively. On the right-hand side, a zoomed-in view of the hydrogen bond network formed by Tau residues and water molecules is displayed. Hydrogen bonds are shown in black, water molecules in pink, and peptide residues in cyan. Figures from (C) was created using Pymol.

**Supplementary Table 1 | X-ray data collection statistics Fyn-SH3/tau(211-220) complex.**

| | |
|---|---|
| Wavelength | 1.0 Å |
| Beamline | SLS-X10SA |
| Detector | EIGER 16M |
| Space group | $P2_12_12_1$ |
| *a* | 35.63 Å |
| *b* | 54.16 Å |
| *c* | 66.87 Å |
| $\alpha, \beta, \gamma$ | 90° |
| MOL (AU) | 2 |
| Resolution[a] | 42.09-1.42 Å (1.47-1.42 Å) |
| No. total reflections | 323,438 (30,235) |
| No. unique reflections | 25,040 (2,408) |
| Redundancy | 12.9 (12.6) |
| Completeness(%) | 99.6 (97.3) |
| Mean $I/\sigma(I)$ | 27 (2.5) |
| $CC^{1/2}$ | 0.999 (0.837) |
| $CC^{*}$ | 1.000 (0.948) |
| $R_{meas}$ (%) | 8.3 (136.9) |
| $R_{pim}$ (%) | 2.3 (38.1) |

[a]Values in parentheses are outer-resolution shell.

**Supplementary Table 2 | X-ray structure refinement statistics Fyn-SH3/tau(211-220) complex.**

| | |
|---|---|
| R-factor | 17.76 |
| $R_{free}$[a] | 21.85 |
| Solvent (%) | 35.7 |
| Mean B-value (Å$^2$) | |
| Protein (main/side) | |
| chain A | 16.93/21.13 |
| chain B | 16.20/20.01 |
| tau peptide | 17.32/18.28 |
| sodium | 17.44 |
| water | 35.05 |
| No. of protein residues | 123 |
| No. of water residues | 95 |
| No. of sodium atoms | 1 |
| Root mean square deviations from ideal geometry | |
| Bond lengths (Å) | 0.015 |
| Bond angles (°) | 1.93 |
| Ramachandran plot (%) | |
| Favoured | 97.96 |
| Allowed | 2.04 |
| Outliers | 0 |

[a]$R_{\text{free}}$ was determined using 5.0 % of the data
